# The adenovirus E4orf1 protein initiates a feedback loop involving insulin and growth factor receptors, AKT, and NF-κB, leading to abnormal DNA content in infected cells

**DOI:** 10.1101/2025.05.12.653445

**Authors:** Madison Moore, Jiang Kong, Ahlam Akmel, Michael A. Thomas

**Author notes:** Corresponding Author: Michael A. Thomas, Ph.D. Department of Biology, Howard University, 415 College Street, NW., Room 333, Washington, D.C. 20059. Present address: The University of Texas MD Anderson Cancer Center, Department of Stem Cell Transplantation and Cellular Therapy, Houston Tx., USA.

## Abstract

Abnormal DNA levels, such as aneuploidy and polyploidy, can indicate cellular transformation and cancer; however, the mechanisms remain poorly understood. All tumor viruses inherently cause abnormal DNA content in cells due to their oncogenes. During infections, adenovirus (Ad) oncogenes—early region 1A (E1A), early region 4 open reading frame 3 (E4orf3), and E4 open reading frame 1 (E4orf1)—promote the abnormal buildup of cellular DNA. Previous studies have described how E1A and E4orf3 lead infected cells to accumulate abnormal DNA content; however, the role of E4orf1 remains speculative. In this study, we generated cells that express E4orf1 to investigate its role in abnormal DNA content. The E4orf1-expressing cells initially exhibited no increase in DNA content compared to the control group. However, after Ad infection, they displayed higher ploidy levels. To detail how E4orf1 influences ploidy levels in Ad-infected cells, we employed pharmacological agents that target E4orf1 signaling. Our results indicate that E4orf1 enhances signaling from insulin and growth factor receptors to AKT and NF-κB, creating a feedback loop that elevates levels of cellular DNA in Ad-infected cells.

**Author Summary:** The early region 4 open reading frame 1 of adenovirus (E4orf1) is recognized for its ability to initiate signals that convert normal cells into cancerous ones. In the initial stages of cancer, cells exhibit DNA content that exceeds the typical levels seen during the G2 and M phases of the cell cycle. This study demonstrates that E4orf1 can trigger a feedback loop involving EGFR, INSR, IGF1R, AKT, and NF-kB, which is both dependent on and independent of PI3 kinase and leads to the accumulation of abnormal DNA content.

## INTRODUCTION

During the early stages of cancer development, some cells increase their genomic content beyond 4N, which is present in G2 and M phase cells [1–4]. The factors that drive this increase in DNA content or polyploidization remain unclear. Adenovirus (Ad), like most tumor viruses [5–8], can promote abnormal cellular DNA content [8, 9]. The main contributors, the Ad oncogenes, include early region 1A (E1A), early region 4 (E4) open reading frame 3 (E4orf3), and early region 4 open reading frame 1 (E4orf1). Previous reports describe how E1A [10] and E4orf3 [9] trigger abnormal DNA levels in cells. However, the mechanism by which E4orf1 contributes has yet to be elucidated.

E4orf1 is an adapter molecule that contains a domain 2 (D2) and a PDZ (<u>P</u>ostsynaptic density protein of 95 kDa (PSD95), <u>D</u>rosophila disc large (Dlg), and <u>Z</u>onula occludens-1 (Zo-1)) protein domain-binding motif (PBM) [11]. These features allow E4orf1 to interact with the epidermal growth factor receptor (EGFR), insulin receptor (InsR), and insulin-like growth factor 1 receptor (IGF1R) [11] and with phosphatidylinositol 3-kinase (PI3K) [12]. In doing so, E4orf1 regulates mitogen-activated protein kinase/extracellular signal-regulated kinase (MAPK/ERK) signaling and further activates the PI3K/AKT/mTOR pathways [11–13]. The signals mediated by E4orf1 are conserved among human Ads [14, 15], allowing infected cells to survive [16]. In this study, we examined the role of E4orf1 in the abnormal DNA content observed in Ad-infected cells. Our research shows that E4orf1 signals from receptor tyrosine kinases (RTKs)—EGFR, InsR, and IGF1R—initiate a feedback loop involving AKT and NF-κB, leading to abnormal DNA content in cells infected with Ad. This feedback loop may be crucial to how E4orf1 converts normal cells into cancer.

## RESULTS

### Effects of E4orf1 on abnormal DNA content in cancer and near-normal cells

In S1A Fig., we identified the different phases of the cell cycle using propidium iodide (PI) [17, 18]. Similar to findings for other tumor viruses [6], replication-defective Ads (*E1B55K*-deleted *dl*1520 [19] and *E4orf6*-deleted *dl*355* [20] in S1C Fig.) induced levels of abnormal DNA content in cells that far exceeded those of the wild-type (*dl*309 Ad [21, 22] (S1B Fig.)). We also noticed that in wild-type Ads (illustrated here by *in*351 in S1B Fig. and elsewhere with other Ads [23]), the inactivation of *E4orf1* does not impact abnormal DNA content in Ad-infected cells due to E1B55K and E4orf6. Therefore, we deleted the *E4orf1* gene from the *E1B55K*-deleted Ad [24], referring to this *E1B55K-* and *E4orf1-*deleted Ad as *E4orf1(**-**)*. As previously reported, more *E1B55K*-deleted (here termed *E4orf1(**+**))* Ad-infected cells accumulated abnormal DNA content than *E4orf1(**-**)* Ad-infected cells [9]. Thus, E4orf1 is associated with enhanced levels of abnormal DNA content in Ad-infected cells.

To confidently declare E4orf1’s role in the abnormal DNA content, we generated GFP- and E4orf1-expressing A549 and HeLa cells (Fig 1A-D). Immunofluorescence microscopy revealed that the negative control cells expressed GFP, while E4orf1-expressing cells did not (Fig 1A and 1C). The HA-tagged GFP (30Kda) and E4orf1 (19Kda) are further distinguished by size using immunoblotting (Fig 1A and 1C). Of the two cell lines, AKT, the primary target of E4orf1 [12, 25], was observed phosphorylated only in the E4orf1-expressing cells (pAkt S473 in Fig. 1A and 1C). Flow cytometry was used to interrogate the cell cycle DNA profiles of the cells. The mock-infected GFP and E4orf1-expressing cells displayed similarly low levels of abnormal DNA content (Fig 1B and 1D).

**Fig 1.**
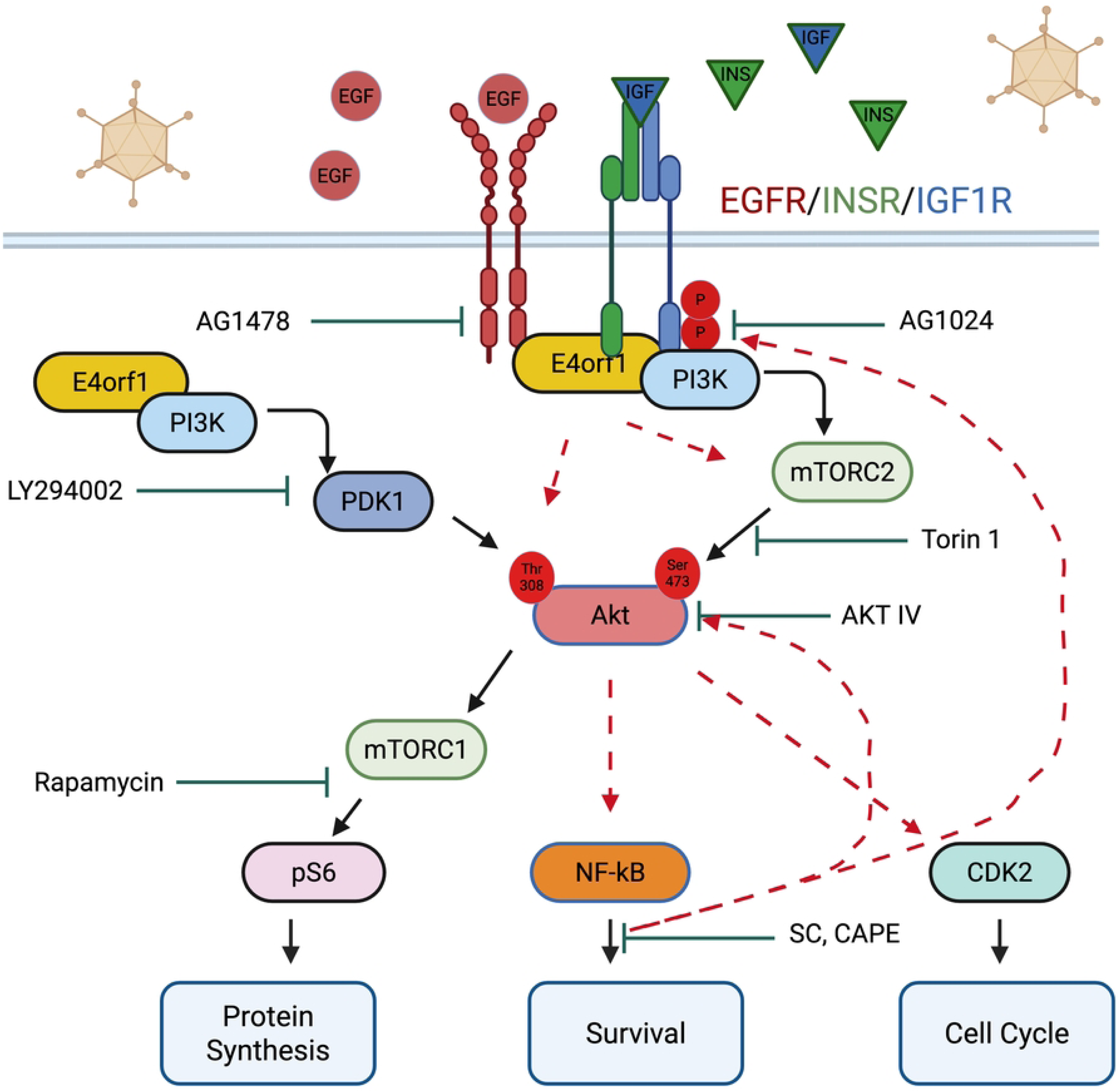
Effects of E4orf1 on the number of cancer and near-normal cells that accumulate abnormal DNA content. (A, B) A549 and (C, D) Hela cells were transfected with MSCVN-HA-GFP or MSCVN-HA-E4orf1 plasmids [27]. The transfected cells were cultured in selection media for 7 to 10 days. (A, C) The cells were photographed at 20X magnification with a spinning disk confocal microscope to confirm the expression of the indicated plasmids. In these images, DAPI stained the nuclei of the cells. (A, C) Immunoblot was used to distinguish the HA-tagged GFP from the HA-tagged E4orf1. Phosphorylated AKT S473 and the Beta (β) tubulin loading control are also shown. (B, D) The HA-tagged GFP-expressing cells and the HA-tagged E4orf1-expressing cells were infected at an MOI of 25 pfu/cell with the indicated Ads for 48 hours. For each group of (B) A549 and (D) Hela cells, the percent cells in each cell cycle phase were plotted in GraphPad prism and are shown. (E) Empty pBabe vector-, Ad9 E4orf1- and Ad5 E4orf1-expressing MCF10A cells were infected with the above-listed Ads and incubated for 48 hours. The *E4orf1(-)* Ad here, *dl*1016 contains the deletion E1B55K, E4orf1, E4orf2 and E4orf3 [28]. Examples of the DNA cell cycle profiles are displayed. The fraction of cells with abnormal DNA content is highlighted in blue.

MCF10A is a non-tumorigenic breast epithelial cell line that provides a reliable model for normal human breast cells [26]. Consequently, we repeated the above experiments in MCF10A cells that stably express either an empty retroviral vector (pBabe) or pBabe containing E4orf1 from Ad subtype 9 (Ad9 E4orf1) or Ad subtype 5 (Ad5 E4orf1) [12, 15]. Similar to the A549 and HeLa cells, only a small fraction of the mock-infected Ad9 and Ad5 E4orf1-expressing MCF10A cells displayed abnormal DNA content (see the last two rows in the first column of Fig 1E). Consequently, the expression of E4orf1 in both cancerous and noncancerous cells does not induce abnormal DNA content.

We then investigated whether expressing E4orf1 in cells could increase the fraction of *E4orf1(**-**)* Ad-infected cells that accumulate abnormal DNA content. More *E4orf1(**-**)* Ad-infected E4orf1-expressing A549, HeLa, and MCF10A cells acquired abnormal DNA content than the mock-infected cells (Fig 1). Thus, unlike when E4orf1 is expressed alone, it enhances the levels of abnormal DNA content that accumulate in both cancerous and near-normal Ad-infected cells.

### Effects of serum depletion on the number of Ad-infected cells with abnormal DNA content

E4orf1 has been reported to promote glucose uptake and increase metabolism in serum-rich and serum-starved environments [29–32]. Therefore, we investigated whether serum levels influenced the ability of Ad to promote abnormal DNA content in cells (Fig 2A and 2D). More *E4orf1(**+**)* Ad-infected cells acquired abnormal DNA content than *E4orf1(**-**)* Ad-infected cells at every serum level evaluated (Fig 2A and 2D). Since lowering the serum levels ultimately reduced the number of cells with abnormal DNA content (Fig 2A and 2D), we conclude that extracellular factors contribute to the propensity for polyploidization in Ad-infected cells.

**Fig 2.**
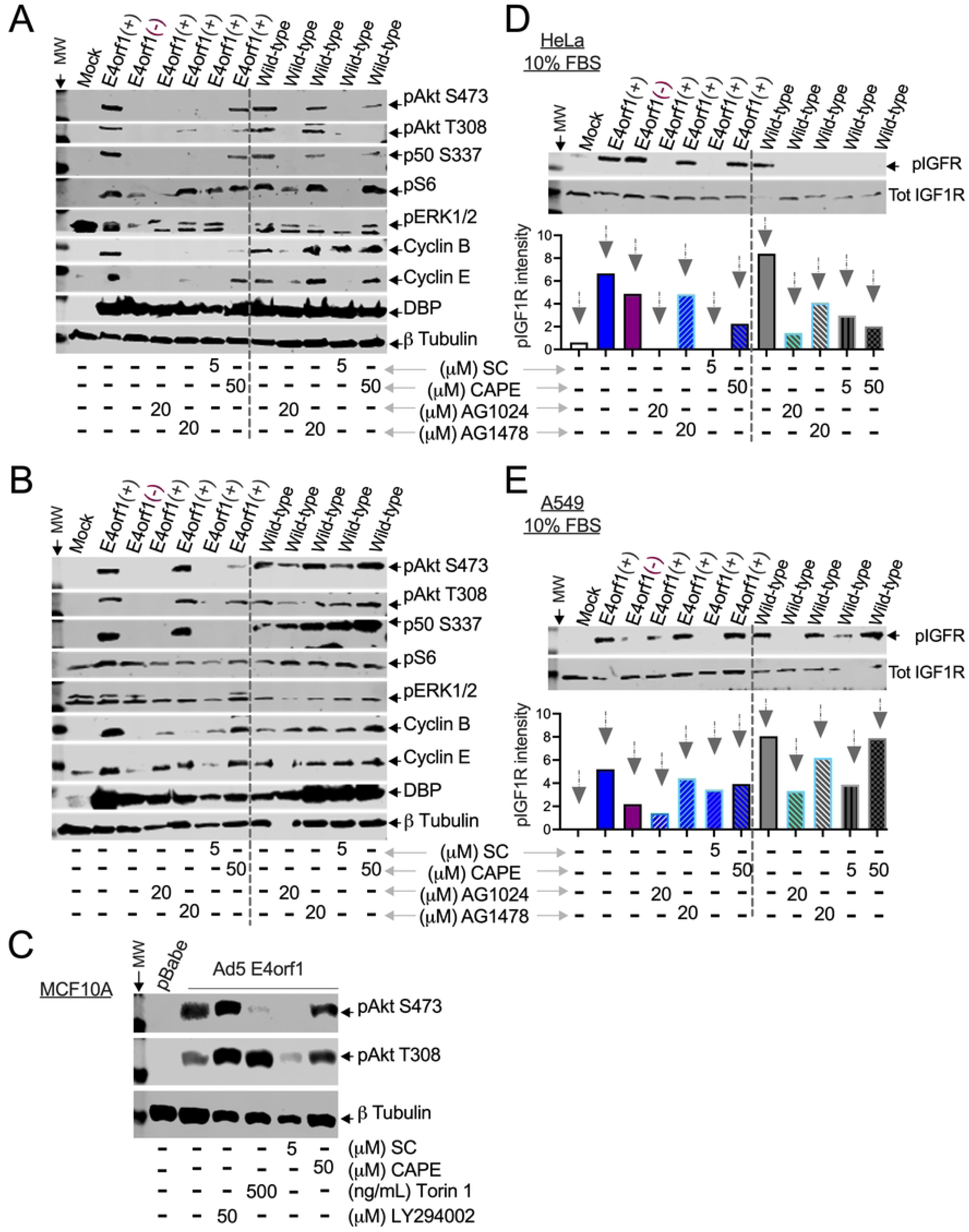
Effects of serum levels on abnormal DNA content in Ad-infected cells. (A) Groups of A549 (n=4-11) and (D) Hela cells (n=4-6) were infected at an MOI of 25 pfu/cell with the *E4orf1(+)* or the *E4orf1(-)* Ad and allowed to incubate in 10%, 1%, or 0% fetal <u>b</u>ovine <u>s</u>erum-containing media (FBS) for 48 hours. The cells were interrogated by flow cytometry. Averages of the percent cells in each phase of the cell cycle with their individual SEM were plotted in GraphPad prism and are shown. The P values were calculated using a two-way analysis of variance (ANOVA) with Holm–Šídák’s multiple comparisons tests. (B, C) A549 and (E, F) HeLa cells were infected with the indicated Ads and incubated for 12, 24, 36, or 48 hours in (B, E) 10% or (C, F) 1% FBS. The analysis of the lysed cells was conducted using immunoblot to identify one or more of the following targets: phosphorylated AKT (pAkt) S473, pAkt T308, NF-κB p50 S337, pS6 S235/S236, pCdk2 T160, cyclins A, B, and E, DBP (Ad infection control), and β-tubulin (loading control).

AKT [33, 34], NF-κB [35, 36], the cyclins, and Cdk2 [37–39] facilitate cell cycle progression and are also known to transmit insulin and growth factor signaling to control cellular metabolism [40–44]. Therefore, we evaluated their status in Ad-infected cells using immunoblots. Regardless of serum levels, a 12-hour incubation with *E4orf1(+)* Ad triggered the phosphorylation of AKT (Fig 2B, 2C, 2E, and 2F). At later times (24, 36, and 48 hpi), in addition to AKT, phosphorylated NF-κB p50, S6, and CDK2 were also detected (Fig 2B, 2C, 2E, and 2F). This starkly distinguished the mock and *E4orf1(-)* Ad-infected cells from those infected with *E4orf1(+)* Ad. Moreover, pharmacological agents that inhibited AKT (AKTIV, Torin 1, LY294002) or its ability to signal to key mediators (Rapamycin) also negatively impacted the levels of abnormal DNA content (S2 and S3 Figs). Consequently, intracellular mediators involved in cell cycle progression and metabolism may affect ploidy levels in Ad-infected cells.

### Effects of Ad on the insulin-like growth factor 1 receptor

E4orf1 interacts with the epidermal growth factor (EGF) receptor (EGFR) and forms a quaternary complex with the insulin (Ins) receptor (InsR) and the insulin-like growth factor 1 (IGF1) receptor (IGF1R) [11] that controls metabolism [29]. Therefore, we evaluated the status of IGF1R in Ad-infected cells grown in 10% FBS over time. At 12 hpi, although IGF1R was detected at similar levels in all the cells, only the *E4orf1(+)* Ad-infected cells showed phosphorylated AKT (Fig 3A). Noticeably, at 24 hpi, levels of IGF1R appeared somewhat reduced in *E4orf1(**+**)* compared to mock and *E4orf1(**-**)* Ad-infected cells (Fig 3A). At 36 and 48 hpi, the IGF1R antibody failed to detect IGF1R in *E4orf1(+)* Ad-infected cells, while it was still detectable in mock and *E4orf1(-)* Ad-infected cells (Fig 3A).

**Fig 3.**
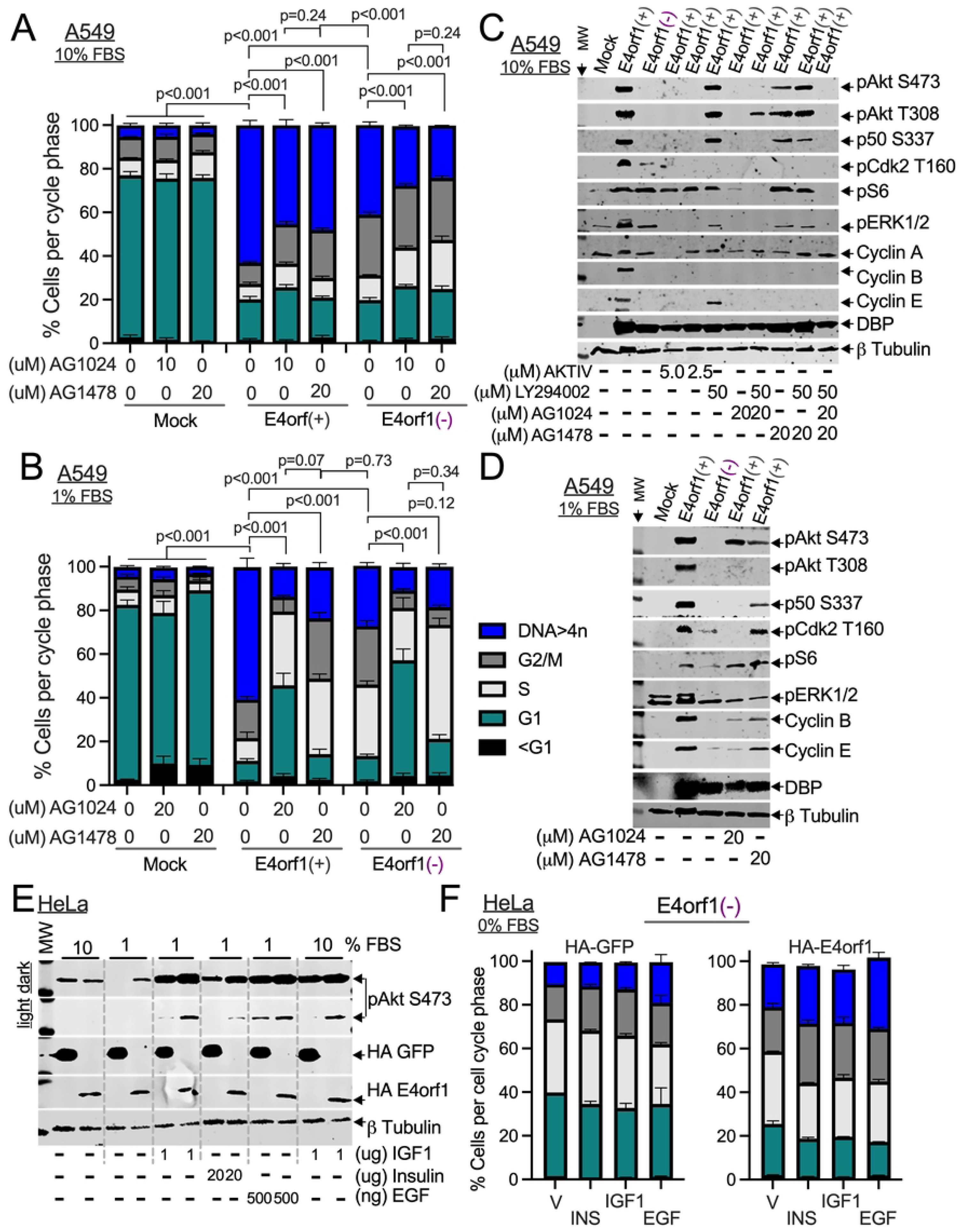
Effects of Ad on the insulin-like growth factor 1 receptors (A) HeLa cells were infected with the indicated Ads for 12, 24, 36, and 48 hours. The cells were lysed and analyzed by immunoblot probing for pAkt S473, IGF1R, and β-tubulin. (B) Hela cells were infected with the indicated Ads, and four hpi were exposed to either AG1478, AG1024, or LY294002 and allowed to incubate in 10% FBS for 48 hours. The lysed cells were analyzed by immunoblot probing for IGF1R, pIGF1R Tyr1135/1136, and β tubulin. The normalized pIGF1R Tyr1135/1136 intensity values were plotted and shown below.

The results in Fig 3A suggest that the IGF1R may be modified during Ad infection. Because phosphorylation of IGF1R has been reported [11], we evaluated this in our systems. Surprisingly, the IGF1R was phosphorylated in both *E4orf1(+)* and *E4orf1(**-**)* Ad-infected cells (Fig 3B). Thus, E4orf1 may not be the only Ad product that stimulates the phosphorylation of the IGF1R during Ad-infection.

Given that the IGF1R levels were again reduced in the *E4orf1(+)* Ad-infected cells, we normalized the levels of phosphorylated IGF1R. According to the normalized values (bar graph at the bottom of Fig 3B), the IGF1R phosphorylation was approximately twice as high in *E4orf1(+)* Ad-infected cells compared to *E4orf1(-)* Ad-infected cells (Fig 3B). Exposure of the *E4orf1(+)* Ad-infected cells to either AG1478, which blocks EGFR activity, or AG1024, which blocks IGF1R activity, reduced the phosphorylation of the IGF1R (Fig 3B). This aligns with other studies [11] and supports an activated EGFR and IGF1R in Ad-infected cells.

### Effects of insulin and growth factor receptors on abnormal DNA content in Ad-infected cells

E4orf1’s interactions with EGFR, InsR, and IGF1R [11] induce metabolic changes [29] that may precede cell transformation [45]. In Fig 3B, we confirmed the mutual dependency of IGF1R on EGFR for activation in Ad-infected cells. Therefore, we evaluated how inhibiting the activity of insulin and growth factor receptors affects abnormal DNA content. Although AG1024 effectively reduced the fraction of Ad-infected cells with abnormal DNA content at a 10μM concentration (Fig 4A), it was more consistent at 20μM, which was used throughout. While AG1024 and AG1478 effectively reduced the number of Ad-infected cells that accumulated abnormal DNA content, at 20μM, AG1024 had a greater impact than AG1478 (Fig 4B, and S4B Fig). Nevertheless, signals from IGF1R and EGFR are essential for Ad-infected cells to accumulate abnormal DNA content.

**Fig. 4:**
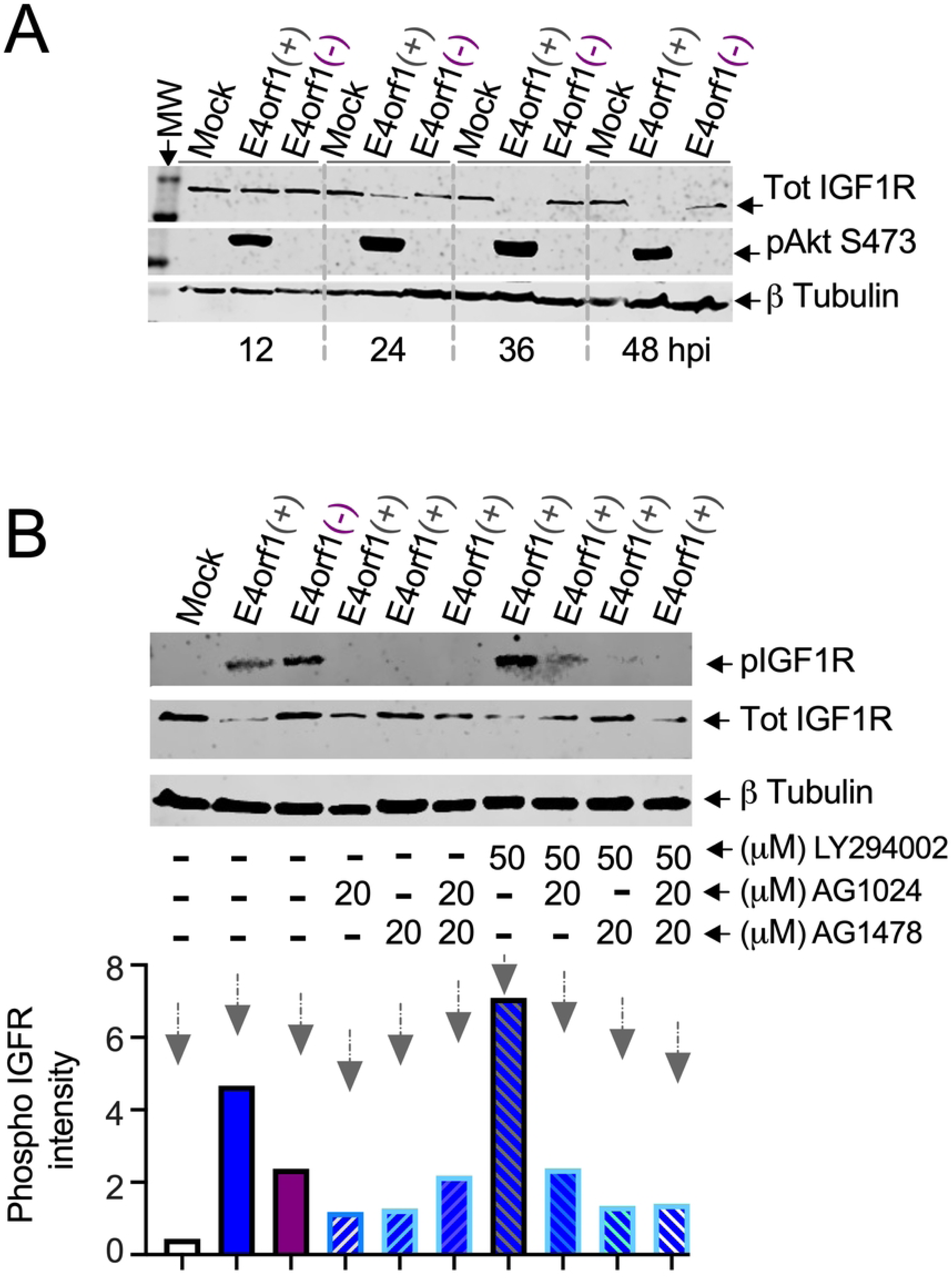
Effects of insulin and growth factor receptors on abnormal DNA content in Ad-infected cells (A, B) A549 cells were infected with the below-indicated Ads, and four hpi were exposed to the indicated concentrations of AG1478 or AG1024. (A) The cells were incubated in 10% or (B) 1% FBS for 48 hours. The stained cells were interrogated by flow cytometry. For (A) n=3-6, (B) n=5-9, and (C) n=3-6, averages of the percent cells in each phase of the cell cycle with their individual SEM were plotted in GraphPad prism and are shown. The P values were calculated using a two-way analysis of variance (ANOVA) with Holm–Šídák’s multiple comparisons tests. (C, D) A549 cells were infected with the indicated Ads. Four hpi, the cells were exposed to either the epithelial growth factor receptor inhibitor AG1478, the insulin growth factor 1 receptor inhibitor AG1024, the AKT inhibitor AKTIV, or the PI3K inhibitor LY294002 at the specified concentrations and allowed to incubate in (C) 10% or (D) 1% FBS for a total of 48 hours. The lysed cells were analyzed by immunoblot probing for a subset of markers including pAkt S473, pAkt T308, NF-κB p50 S337, pS6 S235/S236, pCdk2 T160, pERK1/2 T202/Y204, cyclin A, cyclin B, cyclin E, DBP, and β-tubulin. (E) HA-tagged GFP- and HA-tagged E4orf1-expressing HeLa cells were incubated in 1% FBS for 24 hours. After that, the cells were stimulated with or without 1μg/mL IGF1, 20μg/mL INS, or 500ng/mL EGF and incubated in either 10% or 1% FBS for 30 minutes. As described in the method section, the lysed cells were analyzed by western blot probing for pAkt S473, HA, and β tubulin. (F) HeLa cells expressing HA-tagged GFP or HA-tagged E4orf1 were incubated in 0% FBS and infected with or without the *E4orf1(-)* Ad at an MOI of 25 pfu/cell. Four hpi, the cells were stimulated with 1μg/mL IGF1, 20μg/mL INS, or 500ng/mL EGF. Twenty-four hours later, the stained cells were interrogated by flow cytometry. For each group (n=2), averages of the percent cells in each cell cycle phase with their standard deviations (SD) were plotted in GraphPad prism and are shown.

We investigated the effects of inhibiting EGFR and IGF1R on the phosphorylation of AKT, NF-κB p50, ERK1/2, and CDK2 in *E4orf1(+)* Ad-infected cells. The impact of AKTIV is presented in S2up Fig. It is used here for comparison. As shown, inhibiting IGF1R was as effective at reducing the phosphorylation of AKT, NF-κB p50, ERK1/2, and CDK2 in *E4orf1(+)* Ad-infected cells as AKTIV (Fig 4C). In 10% FBS, LY294002 at the specified concentration can inhibit AKT S473 phosphorylation (S2A Fig). However, this effect is not observed in *E4orf1(+)* Ad-infected cells (Fig 4C, 4D, and S2, S4C, and S4D Figs [16]). Its effectiveness, however, is reflected in the reduced phosphorylation levels of the downstream effectors CDK2 and S6 (Fig 4C, 4D, and S2, S4C, and S4D Figs). Comparatively, AG1478, while not as effective as AG1024, was more effective at inhibiting the phosphorylation of AKT in 10% FBS than LY294002 (Fig 4C, 4D, and S4C, S4D Figs). In 1% FBS, all inhibitors notably reduced the phosphorylation of AKT, ERK1/2, and NF-κB p50 when applied (Fig 4D and S4D Fig). Thus, insulin and growth factor receptors play a vital role in the phosphorylation of AKT, ERK1/2, NF-κB p50, S6, and CDK2 in *E4orf1(+)* Ad-infected cells.

Insulin and growth factor receptor activation is caused by ligand binding [46]. Consequently, we evaluated each ligand’s impact on the phosphorylation of AKT in GFP and E4orf1-expressing cells. The GFP and E4orf1-expressing HeLa cells incubated in 10% FBS exhibited phosphorylated AKT (Fig 4E 1^st^ two columns). By contrast, when the cells were grown in 1% FBS, only the E4orf1-expressing cells exhibited phosphorylated AKT (Fig 4E compare 2^nd^ two columns). Following ligand stimulation, all the cells had increased levels of phosphorylated AKT (Fig 4E see the 3^rd^, 4^th^, 5^th,^ and last two columns). Nonetheless, examining shorter exposure times indicated that AKT exhibited a higher phosphorylation level in cells expressing E4orf1 (see light exposure in Fig 4E, 3rd, 4th, 5th, and last two columns).

Next, we evaluated the individual ligands’ impact on the proportion of Ad-infected cells accumulating abnormal DNA content. The mock-infected GFP- and E4orf1-expressing cells did not show any accumulation of abnormal DNA content, regardless of ligand stimulation (Sup Fig. 4E). However, following infection with the *E4orf1(**-**)* Ad, both cell lines did (Fig 4F, and S4E Fig). Compared to the infected GFP-expressing cells, the infected E4orf1-expressing cells accumulated higher levels of abnormal DNA (Fig 4F and S4E Fig). Furthermore, only the infected E4orf1-expressing cells exhibited increased levels of abnormal DNA content after ligand stimulation (Fig 4F and S4E Fig). In low FBS, E4orf1 induces PI3K-dependent phosphorylation of AKT (S2F Fig). When insulin and growth factor receptors are activated by their ligands, this leads to elevated levels of phosphorylated AKT, indicating that E4orf1-expressing cells respond more effectively to insulin and growth factor receptor signals. The higher the induced AKT, the more Ad-infected cells exhibit abnormal DNA content.

### Effects of NF-κB on the IGF1R and AKT in Ad-infected cells

Inhibiting NF-κB transcriptional activity in Ad-infected cells disrupts their ability to accumulate abnormal DNA content [9]. In Fig 4 and S2 and S3 Figs, we demonstrated that pharmacological agents such as AKTIV, AG1024, and, to a lesser extent, AG1478, which block AKT phosphorylation, also reduced the phosphorylation of NF-κB p50 at S337. The S337 phosphorylation of NF-κB p50 is necessary for its DNA binding activity [47]. This suggests that NF-κB acts downstream of AKT in Ad-infected cells. To further investigate the relationship between NF-κB and AKT, we exposed Ad-infected cells to the NF-κB inhibitors SC and separately to CAPE (Fig 5A and 5B). The effects of AG1024 and AG1478 are presented in Fig. 4 and S4 Fig. and are used here for comparison. As shown, SC was as effective at decreasing AKT phosphorylation in Ad-infected cells as the IGF1R inhibitor (Fig 5A and 5B). CAPE was less effective than SC, likely because of the differences in mode of action. SC blocks NF-κB DNA binding, while CAPE blocks its nuclear translocation. These results show that NF-κB activity is essential for AKT phosphorylation in Ad-infected cancer cells.

**Fig. 5:**
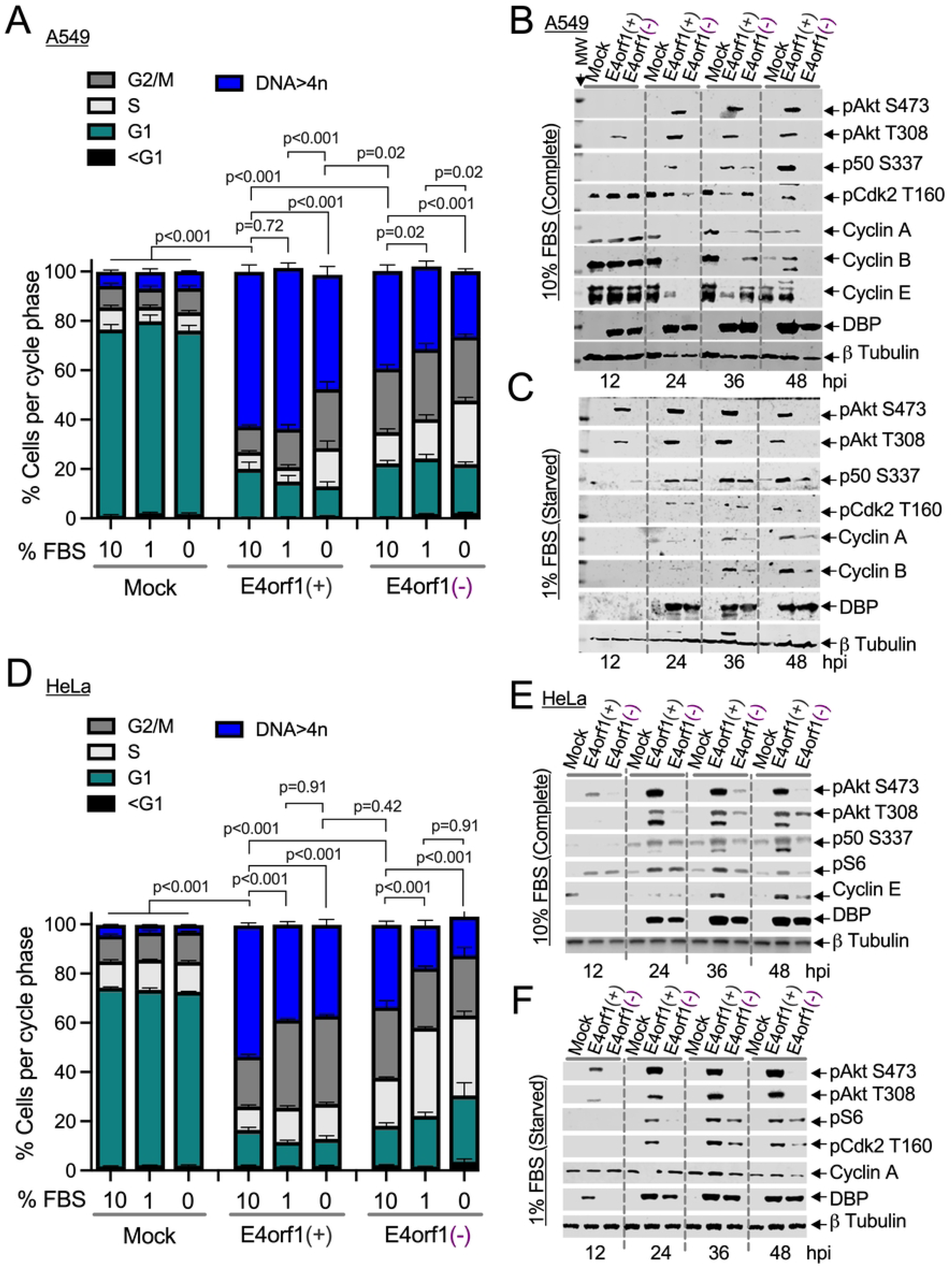
NF-κB mediates the phosphorylation of the IGF1R and AKT in Ad-infected cells. (A) HeLa and (B) A549 cells were infected with the indicated viruses, and four hpi were exposed to AG1024, AG1478, an NF-κB nuclear translocation inhibitor, caffeic acid phenethyl ester (CAPE) [48], or the NF-κB transcriptional inhibitor, SC75741 (SC) [49], at the indicated concentrations for 48 hours. The lysed cells were analyzed by immunoblot probing for pAkt S473, pAkt T308, NF-κB p50 S337, pS6 S235/S236, pCdk2 T160, pERK1/2 T202/Y204, cyclin B, cyclin E, DBP and β-tubulin. (C) Empty pBabe vector- and Ad5 E4orf1-expressing MCF10A cells were exposed either to SC, CAPE, Torin-1, or LY294002, and 48 hours after analyzed by immunoblot probing for either pAkt S473, pAkt T308, or β-tubulin. (D) HeLa and (E) A549 cells were infected with the indicated viruses, and four hpi were exposed to AG1024, AG1478, SC, or CAPE at the indicated concentrations for 48 hours. The lysed cells were analyzed by immunoblot probing for IGF1R, pIGF1R Tyr1135/1136. The normalized pIGF1R intensity values were plotted in GraphPad prism and are shown.

To extend our findings to “normal” cells, we treated MCF10A cells expressing E4orf1 with SC and CAPE, using LY294002 and Torin-1 for comparison. Consistent with results from A549 and HeLa cells (Figs 3, 4, 5A, 5B, and S2 and S4 Figs), LY294002 did not reduce the phosphorylation of AKT in the *E4orf1*-expressing MCF10A cells (Fig 5C). Additionally, as shown in S3 Fig, Torin-1 reduced the phosphorylation of AKT S473 without changing the levels of phosphorylated T308 (Fig 5C). SC reduced the phosphorylation of AKT more effectively than CAPE (Fig 5C). Thus, NF-κB is essential for the phosphorylation of AKT in E4orf1-expressing “normal” cells as well.

We established in Fig 4 that IGF1R activation is crucial for AKT phosphorylation in Ad-infected cells. Therefore, we investigated the effect of NF-κB inhibition on IGF1R phosphorylation. In cells infected with *E4orf1(+)* Ads (both *E1B55K*-deleted on the left and wild-type Ad on the right), IGF1R levels were lower compared to those in the mock and cells infected with *E4orf1(-)* Ad (Fig 5D and 5E). However, after normalization and consistent with Fig. 3, the plotted values for phosphorylated IGF1R were highest in *E4orf1(+)* cells compared to those infected with *E4orf1(-)* Ad (bottom of Fig 5D and 5E). Notably, here, too, SC reduced IGF1R phosphorylation similarly to AG1024, while CAPE produced results comparable to AG1478 (bottom of Fig 5D and 5E, *E1B55K*-deleted on the left and wild-type Ad on the right). Therefore, the pivotal role of NF-κB concerning abnormal DNA content [9] is likely due to its support for the phosphorylation of IGF1R and AKT in Ad-infected cells.

## DISCUSSION

Receptor tyrosine kinases (RTKs), like the EGFR and IGF1R, transmit extracellular signals that control cell cycle progression to include the necessary metabolic processes. Due to gene amplification, overexpression, and mutation, their misfiring can lead to cancer. Abnormal DNA content, such as aneuploidy and polyploidy, are recognized precursors to cancer readily observed in infections with tumor viruses [5, 6, 8]. Ads are prototypical tumor viruses that have been shown to cause cancer [50]. We show here that RTKs can promote abnormal cellular DNA content [51, 52]. In the Ad-infected cell, this is linked to E4orf1.

The individual expression of E4orf1 can stimulate the phosphorylation and activation of AKT. Activating AKT at levels similar to those induced by E4orf1 may be beneficial, as it creates a niche that allows primary cells to survive without serum or cytokines [53]. This ability of E4orf1 can be harnessed to extend the life of CAR T and CAR-NK cells, which are currently being explored as tumor therapy [54, 55]. E4orf1 can also improve metabolism [56–58]. Thus, it is being explored for diabetes treatment [59], for improving cognition in Alzheimer’s disease [60], and for reducing aging markers [61].

In light of our findings, we recast the roles of the Ad oncogenes E1A, E1B, E4orf6, E4orf3, and E4orf1 [62, 63]. E1A’s interactions with pRB and p300 promote unscheduled DNA synthesis, leading to massive cell death [64–67]. Some cells capable of activating ancient polyploidization programs [68, 69] can survive and become transformed [70, 71]. *E1B* and *E4orf6* enhance the number of E1A-expressing cells that transform [62, 63]. However, during infection, *E1B* and *E4orf6* prioritize viral progeny [19, 20, 72] over oncogenic transformation by increasing the availability of cytoplasmic viral RNA [72–75], thus minimizing the accumulation of cells with abnormal DNA content. Like E1B and E4orf6, E4orf3 also enables more E1A-transformed cells to survive [76]. E4orf3, by disrupting MRN complex formation, enhances levels of phosphorylated ATM [9]. Although our idea that ATM is active in Ad-infected cells differs from the prevailing views [77], the notion that ATM might be necessary to fully activate AKT [78] and maintain NF-κB levels [79–81] is backed by evidence suggesting that ATM is needed for IGF1R expression [82, 83] and the phosphorylation/activation of AKT and NF-κB in Ad-infected cells [9]. This connects E4orf3 to E4orf1, clarifying why we assert that in Ad-infected cells, E4orf3 collaborates with E4orf1 [9].

NF-κB has been reported to act downstream of AKT [84–86]. However, our study also suggests that NF-κB may function upstream of AKT (Fig 6). While this concept is new for Ad, the idea that NF-κB can act upstream of AKT may not be novel. In one study, overexpression of NF-κB resulted in increased AKT mRNA and protein expression [87]. In another, a positive feedback loop involving EGFR/AKT/mTORC1 and IKK/NF-κB was shown to influence proliferation in head and neck squamous cell carcinoma [88]. E4orf1, therefore, due to its capacity to integrate signals from various RTKs, may help maintain this AKT-NF-κB feedback loop, allowing polyploidization and transforming normal cells into cancer cells [14, 89]. While the signals we discussed may be necessary for E4orf1 to function as an oncogene, it is important to note that abnormal DNA content (aneuploidy and polyploidy) is also linked to aging and neurodegenerative diseases [90–93]. The signals outlined here could also be significant there.

**Fig 6.**
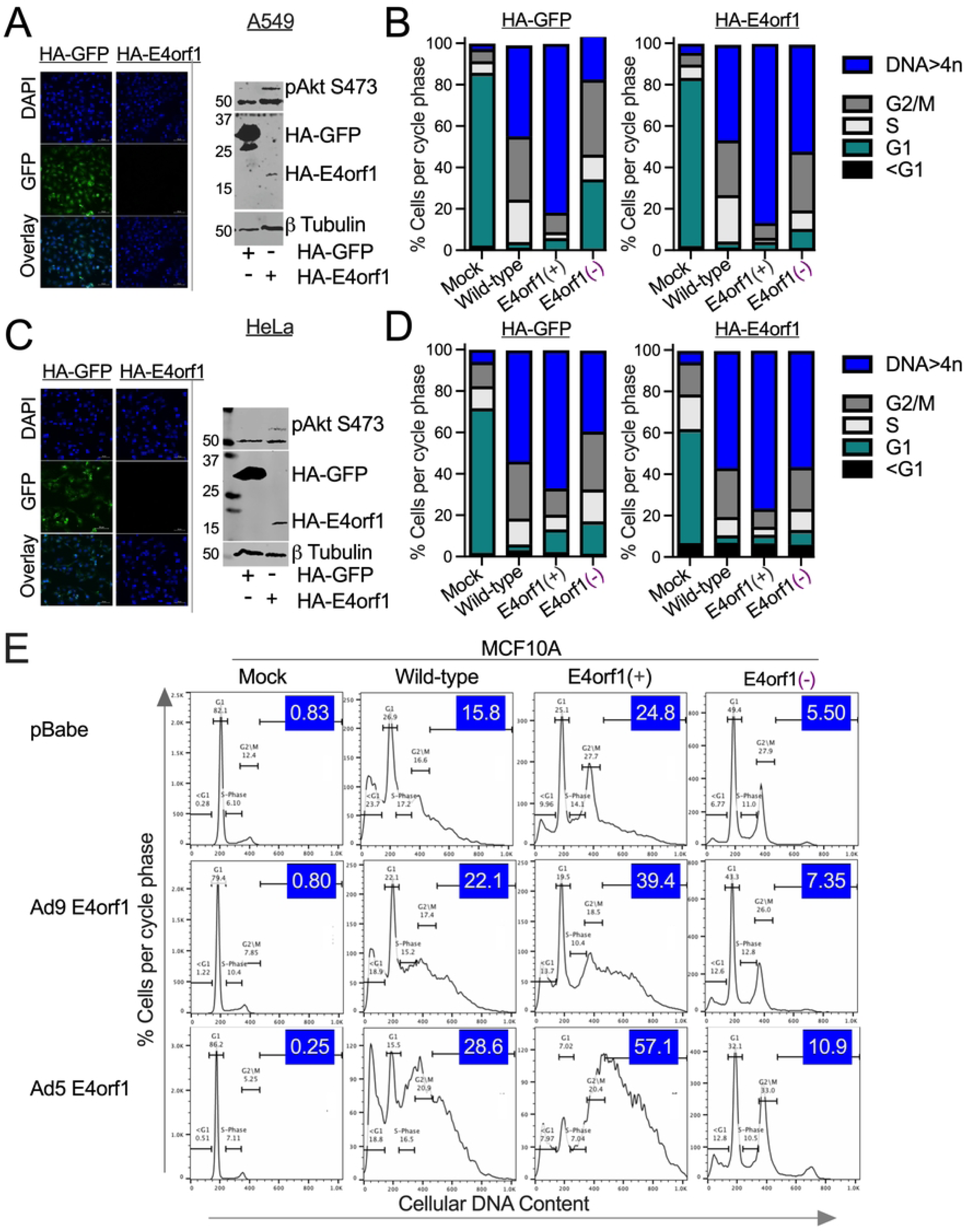
Schematic illustrating E4orf1-mediated signals that promote aneuploidy and polyploidy. In the presence of EGF, INS, and IGF1, and during Ad infection, E4orf1 activates a [PI3 kinase-dependent and independent] feedback loop involving EGFR, INSR, IGF1R, AKT, and NF-kB that enhances essential cellular activities, such as protein synthesis, cell survival, and progression through the cell cycle, all of which are vital for accumulating abnormal DNA content. (This image was created using BioRender)

## Author Contributions

Conceptualization, M.A.T; Methodology, M.A.T.; Formal Analysis, M.M., and M.A.T.; Investigation, A.A., J.K., M.M., Resources, M.A.T.; Writing – Original Draft, M.M., and M.A.T.; Writing –Review & Editing, A.A., J.K., M.M., M.A.T.; Visualization, M.M., and M.A.T.; Supervision, M.A.T.; Project Administration, M.A.T.; Funding Acquisition, M.A.T.

## Methods section

### Contact for reagent and resource sharing

Further information and requests for resources and reagents should be directed to

### Ethics Statement

All experiments were approved by the Institutional Biosafety Committee (IBC) at Howard University.

### Cell culture

We used the human cervical carcinoma-derived HeLa cell line (ATCC^®^ CCL-2^™^) because most of what is known about Ad is in the context of this cell line. We used the human lung carcinoma epithelial A549 cell line, A549 (ATCC^®^ CCL-185), as Ad is known to target the upper respiratory airway. The HeLa and A549 cell lines were cultured in Dulbecco’s Modified Eagle *Medium* supplemented with 10, 1, or 0% Fetal Bovine Serum (FBS), 100 unit/ml Penicillin, and 100 mg/ml Streptomycin. The human breast mammary epithelial cell line MCF10A stably expressing pBabe-empty or pBabe-E4orf1 were gifts received from Ronald T. Javier, Baylor College of Medicine, Houston, Texas, and were described previously [12, 15]. MCF10A cells were cultured with DMEM/F12 medium supplemented with 5% horse serum (Life Technologies, Carlsbad, CA) and MEGM^TM^ Mammary Epithelial Cell Growth Medium SingleQuots^TM^ Kit (Lonzo Cat # CC-4136). All cells were incubated at 37°C in humidified air under 5% C0_2_.

### Viruses

The adenovirus subtype 5 (Ad5) hr404 and the E1B55K- and *E4orf1*-deleted *Δ*E4orf1 Ad have been described before [23]. The Ad5 *dl*309 is another commonly used phenotypic Ad wild type that contains a deletion in the early region 3 (E3) [21, 22]. This Ad has been used to create most of the Ad vectors used clinically. Ad5 in351 includes a 5-bp insertion within the E4orf1 coding region that inactivates the E4orf1 [72]. The mutant Ad5 *dl*1520 (listed here as *ΔE1B*) has an 827-bp deletion in the region encoding the 55kDa protein [19]. The mutant Ad5 *dl3*55* contains a 14-bp deletion that inactivates E4orf6 [20].

### Infection

Cells were plated in 6-well plates at a density of 5 x 10^5^ cells per well or in 12-well plates at a density of 1.5 x 10^5^ cells per well and infected at an MOI of 25 with the indicated Ads by incubating for 1 hour at 37°C while rocking every 10-15 minutes. The media was aspirated and replaced with 2ml (6-well dish) or 1ml (12-well dish) of fresh medium and incubated for the specified times.

### Flow Cytometry

Cells were detached from the plate using trypsin, washed in 1x PBS, and fixed in 70% ethanol overnight at −20°C. The fixed cells were then washed twice with PBS. After centrifuging, the pellet was stained with 500ul of FxCycle^TM^ PI/RNase staining solution (ThermoFisher Cat #: F10797) per 1 million cells. The samples were incubated at room temperature for 15–30 minutes, protected from light. The cells were acquired with a BD (Becton, Dickinson) FACSVerse™ flow cytometer. The population of cells in each phase of the cell cycle was quantified using FlowJo™ v10.8 Software (BD Life Sciences), as shown in Supplemental Fig. 1A.

### Western Blot

Cells were lysed in 1X SDS sample buffer (ThermoFisher Cat #: LC2675-4) containing protease/phosphatase inhibitor (ThermoFisher Cat #:78442) and 5% Beta-mercaptoethanol (BME). Equal amounts of lysate were loaded into wells of 4-20% Tris-Glycine gels (ThermoFisher Cat #: XP04205BOX). Following electrophoresis, proteins were transferred from the gel to a nitrocellulose membrane (ThermoFisher Cat #: IB23002). At room temperature, the membrane was blocked in PBS with 0.02% Tween-20 and 20% milk for one h. After blocking, the membrane was incubated in appropriate primary antibody dilutions in PBS with 0.02% Tween-20 and 10% milk on a rocker at four °C overnight. After washing three times in PBS with 0.02% Tween-20 and 5% milk for 5 min each time, the membrane was incubated with appropriate secondary antibody dilutions in PBS with 0.02% Tween-20 and 10% milk for 30 minutes at room temperature. The membrane was washed three times, 5 min each, and exposed to a 1:1 substrate luminal/enhancer solution, and the image was captured using a LI-COR Odyssey® Fc Imaging System (Lincoln, Nebraska, USA).

In each case, the blots were obtained by probing for one antigen, stripping with Restore™ Western Blot Stripping Buffer (ThermoFisher Cat #:21059), and probing again for another antigen after washing and blocking as above. Where indicated, the Ad <u>D</u>NA <u>b</u>inding <u>p</u>rotein (DBP) was used as an infection control and Beta (β) tubulin as a loading control. The lane denoting the molecular weight marker, MW, is on most blots.

### Western Blot Normalization

Fluorescent intensity was measured using the LI-Cor Image Studio Lite program. Each blotted area was manually gated, and intensity values were transferred to Excel. The lane normalization factor was determined by dividing each lane’s observed IGF1R signal intensity by the highest IGF1R fluorescent value on the membrane. The pIGF1R signals were divided by the lane normalization value. The normalized values were plotted using GraphPad prism.

### Antibodies

The antibodies used for western blot and the dilutions used were Akt ser473 (Invitrogen Cat #: 44-62G), 1:1000; Akt thr308 (Cell signaling Clone #: D25E6), 1:1000; Beta Tubulin (Invitrogen Cat #: PA1-16947), 1:5000; Cyclin A (BD Cat #: 611268), 1:1000; Cdk2 T160 (Santa Cruz Biotech Cat #: SC-101656), 1:250; cyclin E (BD Cat:51-1459GR), 1:1000; IGF1R (Invitrogen Cat #: PA5-85986), 1:1000; HA Tag (Invitrogen Cat #: 26183), 1:5000; p50ser337 (Santa Cruz Biotech Cat #: SC-271908), 1:1000; p-S6 Ribosomal Protein S240/244 (Cell signaling Clone #: D68F8), 1:5000; P-p44/42 MapK T202/Y204 (cell signaling Clone #: D13.14.4E), 1:5000; cyclin B (BD Cat #: 610219), 1:1000; and P-IGF-1R beta Y1135/1136/InsR beta (Cell signaling Clone #: 19H7), 1:1000. The western blot marker used was the LI-COR molecular weight marker 928-40000.

### Pharmacological Inhibitors

The pharmacological inhibitors used in this study were Torin1 (Cayman Chemical Company Cat #: 10997), LY294002 (Selleckchem Cat #: S1105), Akt inhibitor IV (Millipore Cat #: 124011), SC75741 (Selleckchem Cat #: S7273), AG-1478 (Selleckchem Cat #: S2728), AG-1024 (Selleckchem Cat #: S1234), Caffeic Acid Phenethyl Ester (CAPE) (Selleckchem Cat #: 7414), and rapamycin (Selleckchem Cat #: S1039).

### Plasmid Transfection

The plasmids MSCV-N E4orf1 (Addgene plasmid # 38063) and MSCV-N GFP (Addgene plasmid # 37855) were gifts from Karl Munger and were described before [27]. Bacterial stabs were streaked on LB agar plates (Quality Biological Cat #: 340-1070-231) containing 50μg/ml of ampicillin, which were then incubated for 12 hours at 37°C. Following incubation, individual colonies were picked using a sterile tip and placed in a 50 ml conical tube containing 25 ml of LB broth (KD Medical Cat #: BLF-7030), supplemented with 100mg/ml of ampicillin. The tube was then placed in a shaker for 12 hours at 250rpm at 37°C for bacterial amplification. Following shaker incubation, plasmid DNA was isolated using the Pure Link HiPure Plasmid Filter Maxiprep Kit (ThermoFisher Cat #: K210017), yielding pure plasmid DNA. The purified DNA was then transfected into cells using lipofectamine 2000. After 24 hours, transfected cells were incubated in 1um of puromycin for selection. Immunofluorescence microscopy was used to verify GFP expression, and a western blot was used to verify the expression of HA-tagged GFP and HA-tagged E4orf1.

### Immunofluorescence

Cells were seeded on coverslips at a density of 2.5 × 10^5^ cells/well in 12-well plates and incubated at 37°C for 24 hours. Cells were washed twice with 1x PBS and fixed with 4% paraformaldehyde for 15 minutes at room temperature. Subsequently, cells were permeabilized in 0.5% TritonX-100 (PBS) for 10 minutes at room temperature. The cells were rinsed with 1x PBS and stained with DAPI Fluoromount-G (Invitrogen Cat #: 00-4959-52). Images were obtained using a Nikon Ti-E-PFS inverted microscope with a 100× 1.4 NA Plan Apo Lambda objective. The microscope also had a Yokogawa CSU-X1 spinning disk unit and an Andor iXon 897 EMCCD camera. To analyze the DNA and GFP in both HeLa and A549 cells, excitations were selected at 405 nm and 488 nm, respectively. All images are overlay z-plates.

### Statistical analysis

Two-way analysis of variance (ANOVA) was used with Holm-Šídák’s multiple comparisons test to compare mean differences in cell cycle phases between the groups. P-values < 0.05 are considered significant.

## Acknowledgments

We thank Dr. David Ornelles (Wake Forest University) for the *dl*1520 and *dl3112* virus, DBP hybridoma cells, and Ronald T. Javier (Baylor College of Medicine, Husto) for the MCF10A stably expressing pBabe-empty and pBabe-E4orf1 cell lines. We also thank Dr. Sergei Nekhai (Howard University) for access to the BD FACSVerse, and Dr. Anna Allen (Howard University) for access to the Spinning Disk Confocal Microscope.

## Supporting information

**S1 Fig. Effects of Ad on the number of infected cells with abnormal DNA content.**

**(A)** HeLa cells were mock- or infected at a multiplicity of 30 plaque-forming units per cell (MOI30 pfu/cell) with an *E1B55K*-deleted Ad (*ΔE1B*) or an *E1B55K*- and E4orf1-deleted Ad (*ΔE1B/E4orf1*). At 48 hpi, the cells were washed, stained in an RNase containing propidium iodide solution, and analyzed using flow cytometry. Propidium iodide (PI) binds to cellular DNA, which allows for determining different cell cycle phases [20, 21]. (**i**) First, we gated the cells to remove doublets and other clumps that can confound the results. Cells with fragmented DNA or dead cells have DNA content less than the 2N (where N is the chromosome number) found in G1 phase cells (this fraction is labeled <G1 in **ii-vi**). (**ii, iii**) Most cells growing in culture are in the G1 phase of the cell cycle (here 67.8%). During the S phase, DNA synthesis occurs, leading to 4N DNA content in G2 phase cells. Before cytokinesis, the 4N cells go through the M phase of the cell cycle. The G2 and M phase cells (14.9%) contain similar DNA content and cannot be distinguished by this method. (**iii**) The percentage of cells in each cell cycle phase is indicated. (**iv-v**) By 48 <u>h</u>ours <u>p</u>ost-infection (hpi), 61 percent of the *ΔE1B* Ad-infected cells exhibited abnormal DNA content, compared to 32 percent of the *ΔE1B/E4orf1* Ad-infected cells. (**vi**) An overlay of the various histograms **ii-v** is shown.

**(B)** Deletion of E4orf1 from wild-type Ad has little effect on DNA content. Hela cells were infected at an MOI of 50 pfu/cell with **(top)** wild-type Ad *dl*309 or **(bottom)** *in*351 with an insertion mutation in the *E4orf1* gene for 12 and 24 hours. The stained cells were interrogated by flow as above in A. The percentage of cells with abnormal DNA content for each virus infection is shown in blue. **(C)** More cells infected with E1B55K- and E4orf6-deleted Ads have abnormal DNA content compared to wild-type Ad-infected cells. Hela cells were infected at an MOI of 50 pfu/cell with **(top)** the *E1B55K*-deleted Ad*, dl*1520, or **(bottom)** an *E4orf6*-deleted Ad (*ΔEorf6*) *dl*355* for 12 and 24 hours. The stained cells were treated and analyzed as described in part A.

**S2 Fig. PI3K and AKT signals are needed for cell cycle transition and the accumulation of abnormal DNA content in Ad-infected cells.**

(A) A549 cells were incubated in 1% FBS for 24 hours and then stimulated with 1μg/mL of insulin growth factor 1 (IGF1) for 30 minutes with or without the 50uM PI3K inhibitor LY294002. The vehicle control (v cont) DMSO was used as a negative control. The lysed cells were analyzed by immunoblot probing for pAkt S473, pS6 S235/S236, and β-tubulin.

(B, D) A549 and (C, E) Hela cells were infected at an MOI of 25pfu/cell with the indicated Ads and 4 hours post-infection (hpi) exposed to either 50uM LY294002 or 5uM AKTIV and allowed to incubate in 10% FBS for a total of 48 hours. (B, C) The lysed cells were analyzed for one or more of the following targets using immunoblotting: phosphorylated AKT (pAkt) S473, pAkt T308, NF-κB p50 S337, pS6 S235/S236, pCdk2 T160, pERK1/2 T202/Y204, as well as cyclins A, B, and E, DBP, and β-tubulin.

(D) For each group of A549 (n=3-8) and (E) HeLa cells (n=3-6), averages of the percent cells in each phase of the cell cycle with their individual SEM were plotted in GraphPad prism and are shown. The P values were calculated using a two-way analysis of variance (ANOVA) with Holm–Šídák’s multiple comparisons tests.

(F, G) A549 cells were infected with the indicated viruses, and four hpi were exposed to either LY294002 or AKTIV at the indicated concentrations and allowed to incubate in 10% or 1% FBS for 48 hours. (F) The lysed cells were analyzed by immunoblot probing for pAkt S473, pAkt T308, NF-κB p50 S337, pCdk2 T160, cyclin A, cyclin B, cyclin E, DBP, and β-tubulin.

(G) For each group (n=3-8), averages of the percent cells in each phase of the cell cycle with their individual SEM were plotted in GraphPad prism and are shown. The P values were calculated using a two-way analysis of variance (ANOVA) with Holm–Šídák’s multiple comparisons test.

**S3 Fig. The mTOR complex 1 and 2 mediate abnormal DNA content in Ad-infected cells**

(A) A549 cells were incubated in 1% FBS for 24 hours, followed by stimulation with 1 μg/mL of insulin growth factor 1 (IGF1) for 30 minutes, with or without 50 nM rapamycin or 500 ng/mL Torin1. DMSO served as a negative control. The lysed cells were analyzed by immunoblotting for pAkt S473, pS6 S235/S236, and β-tubulin.

(B, C) A549 cells were infected with the specified viruses. Four hours post-infection (hpi), they were treated with either 50 nM rapamycin or 500 ng/mL Torin1 and allowed to incubate in (B) 10% or (C) 1% FBS for a total of 48 hours. The stained cells were analyzed by flow cytometry. For each group incubated in 10% (n=4-6) and 1% FBS (n=3-8), the average percentages of cells in each phase of the cell cycle, along with their respective standard error of the mean (SEM), were plotted using GraphPad Prism and are presented here. The P values were calculated using a two-way analysis of variance (ANOVA) with Holm–Šídák’s multiple comparisons test.

(D, E) A549 cells were infected with the specified viruses, and four hours post-infection (hpi), they were exposed to either rapamycin or Torin1 at the indicated concentrations, followed by incubation in (D) 10% or (E) 1% FBS for 48 hours. The lysed cells were analyzed using immunoblotting to detect pAkt S473, pAkt T308, NF-κB p50 S337, pCdk2 T160, pS6 S235/S236, cyclin A, cyclin B, cyclin E, DBP, and β-tubulin.

**S4 Fig. Effect of insulin and growth factor receptors on abnormal DNA content in Ad-infected cells**

(A) A549 cells were incubated in 10% FBS for 24 hours and then stimulated with 500ng/mL of epithelial growth factor (EGF) for 30 minutes with or without 20uM of the epithelial growth factor receptor inhibitor AG1478. The lysed cells were analyzed by immunoblot probing for pAkt S473, pERK1/2 T202/Y204, pS6 S235/S236, and β-tubulin.

(B) HeLa cells were infected with the below-indicated Ads, and four hpi were exposed to the indicated concentrations of AG1478 or AG1024 and incubated in 10% FBS for 48 hours. The stained cells were interrogated by flow cytometry. The averages (n=5-9) of the percent cells in each phase of the cell cycle with their individual SEM were plotted in GraphPad prism and are shown. The P values were calculated using a two-way analysis of variance (ANOVA) with Holm–Šídák’s multiple comparisons tests.

(C, D) Hela cells were infected with the indicated Ads. Four hpi, the cells were exposed to either the epithelial growth factor receptor inhibitor AG1478, the insulin growth factor 1 receptor inhibitor AG1024, the AKT inhibitor AKTIV, or the PI3K inhibitor LY294002 at the indicated concentrations and allowed to incubate in (C) 10% or (D) 1% FBS for a total of 48 hours. The lysed cells were analyzed by immunoblot probing for pAkt S473, NF-κB p50 S337, pS6 S235/S236, cyclin A, cyclin B, cyclin E, DBP, and β-tubulin.

(E) A549 cells expressing HA-tagged GFP or HA-tagged E4orf1 were incubated in 0% FBS and infected with or without the *E4orf1(-)* Ad at an MOI of 25 pfu/cell. Four hpi, the cells were stimulated with 1μg/mL IGF1, 20μg/mL INS, or 500ng/mL EGF. Twenty-four hours later, the stained cells were interrogated by flow cytometry. For each group (n=2), averages of the percent cells in each cell cycle phase with their standard deviations (SD) were plotted in GraphPad prism and are shown.

